# Gallium induces cytotoxicity through disruption of DNA synthesis rather than ferroptosis

**DOI:** 10.64898/2026.04.01.715544

**Authors:** Jingzhi Fan, Annija Vaska, Xiping Jiang, Kristaps Klavins

## Abstract

**Background:** Gallium (Ga) is a promising anti-tumor agent; however, its precise molecular targets in osteosarcoma remain debated. While current paradigms largely attribute its toxicity to reactive oxygen species (ROS) and ferroptosis, understanding its true mechanism is essential for overcoming therapeutic resistance. This highlights the need for interdisciplinary approaches, such as metabolomics, to unveil novel vulnerabilities in cancer metabolism.

**Methods:** We employed an interdisciplinary strategy utilizing high-resolution liquid chromatography-mass spectrometry (LC-MS) metabolomics and ^13^C_2_-glutamine stable isotope tracing in osteosarcoma cells to elucidate the cytotoxic mechanisms of gallium nitrate. Scanning electron microscopy with energy-dispersive X-ray spectroscopy (SEM-EDS) was utilized for elemental mapping, and in silico modeling was applied to evaluated metal binding dynamics. Furthermore, synergistic effects were tested by combining gallium with the DNA-damaging agent cisplatin.

**Results:** Our metabolic profiling revealed a profound bifurcation characterized by the systemic depletion of glycolysis and pentose phosphate pathway intermediates, coupled with a novel ribonucleotide accumulation bottleneck. The observed distinct signature strongly implicated ribonucleotide reductase (RNR) as the primary enzymatic target. In silico modeling and SEM-EDS visually and thermodynamically confirmedthat gallium acts as a structural decoy for iron within the RNR active site. The co-localization induces functional iron starvation rather than canonical ferroptosis. Furthermore, isotope tracing confirmed that elevated ROS is a consequence of overall metabolic failure, not the primary driver of cell death. Crucially, gallium functioned as a metabolic DNA repair inhibitor, synergizing potently with cisplatin to prevent the repair of platinum-induced DNA lesions.

**Conclusions:** Gallium selectively sensitizes highly proliferative sarcoma cells by disrupting RNR-mediated DNA precursor synthesis, while sparing normal osteoblasts. Leveraging metabolomics to uncover this state of functional iron starvation provides a rational, interdisciplinary framework for developing gallium-based combination therapies designed to break platinum resistance in clinical oncology.

## Main

Gallium has historically been utilized in medicine for its antibacterial properties and has recently emerged as a promising candidate for cancer therapy, demonstrating substantial anti-tumor activity across various malignancies [1–5]. Clinically, gallium-based compounds, such as KP46, have undergone testing for osteosarcoma [6] while gallium nitrate is currently utilized in the treatment of non-Hodgkin’s lymphoma [7]. Despite its growing translational potential, the fundamental mechanisms underlying gallium-induced cytotoxicity remain obscured by contradictory theories. For instance, a gallium complex featuring planar salen ligands has been reported to trigger ferroptosis [8] and numerous studies attribute gallium’s cytotoxic effects primarily to reactive oxygen species (ROS) and related ferroptotic pathways [9,10]. Conversely, other investigations indicate that gallium triggers caspase-mediated apoptosis [11–13] induces cell-cycle perturbations [14], or specifically suppresses ribonucleotide reductase (RNR) and catalase activity without affecting aconitase [15].

A prevailing hypothesis is that gallium exerts its anti-cancer effects by disrupting iron metabolism. As a group III metal, gallium shares striking physicochemical properties with iron (table S1.); for example, the octahedral ionic radius for Ga^3+^ (0.620 Å) closely mimics that of high-spin Fe^3+^ (0.645 Å) [16]. Because iron is an essential element for cancer cell proliferation and survival, gallium can competitively bind to transferrin receptors, inhibiting receptor-mediated iron uptake. The interference with iron homeostasis subsequently disrupts RNR function, mitochondrial activity, and the regulation of transferrin receptors and ferritin. Furthermore, gallium nitrate has been shown to stimulate an increase in mitochondrial ROS, triggering the downstream upregulation of metallothionein and hemoxygenase-1 [17]. Ultimately, these combined disruptions dismantle the cellular processes required for malignant proliferation. In clinical and research settings, gallium is typically administered as gallium nitrate or gallium citrate, which are approved for treating cancer-related hypercalcemia. Compared to various complexed forms, gallium nitrate is a highly ionized salt [18]. The simple formulation eliminates the confounding variables and synergistic interferences often introduced by complex ligands, making it the optimal candidate for studying the direct cellular effects of gallium ions. Consequently, gallium nitrate itself has been the subject of numerous clinical trials (e.g., NCT00002543, NCT00054808, NCT00002578, NCT00836173).

To fully understand the mechanism of action for gallium and aid the development of next-generation cancer treatment approaches, we utilized an in vitro osteosarcoma model treated with gallium nitrate. By deploying an interdisciplinary approach that integrates metabolomics, stable isotope tracing, and pioneering SEM-EDS elemental mapping, we aimed to map the precise metabolic bottleneck induced by gallium and visualize the competitive distribution of gallium and iron. In this context, advanced metabolomics serves as a critical bridge between analytical chemistry and tumor biology. Furthermore, we utilized ferroptosis inhibitors to delineate whether ROS accumulation acts as a primary driver or a secondary consequence of cell death. Ultimately, understanding these specific molecular targets provides crucial insights for leveraging gallium as a metabolic DNA repair inhibitor, offering a rational foundation for synergistic combination therapies aimed at breaking chemoresistance of platinum through synergistic effects.

## Results

### Distinct metabolic profiles between the two types of bone cells at optimized concentration

, We assessed cell viability across a concentration gradient (0–2000 μM) in normal and malignant bone cell lines over 24 and 72 hours (Fig. 1A) to evaluate the therapeutic window and verify the cytotoxicity of gallium (Ga). All cell lines maintained high viability at low concentrations. However, the results demonstrated a profound selective toxicity: while normal MC3T3-E1 pre-osteoblasts and NIH/3T3 fibroblasts maintained robust viability in the short term, the osteosarcoma cell lines (MG-63 and K7M2) exhibited a significant, dose- and time-dependent reduction in viability. Specifically, MC3T3-E1 cells only displayed decreased viability at very high concentrations (>1000 μM at 24 hours, and >500 μM at 72 hours). Conversely, at 250 μM in the 72-hour assay, gallium exhibited substantial cytotoxicity toward the sarcoma cells, yielding an IC50 markedly lower than that observed for the osteoblasts (Fig. S1C, D). Notably, by 72 hours, the NIH/3T3 and K7M2 cells showed similar cell death gradients. Overall, gallium ions demonstrated selective cytotoxicity against osteosarcoma cells at optimized concentrations.

**Fig. 1:**
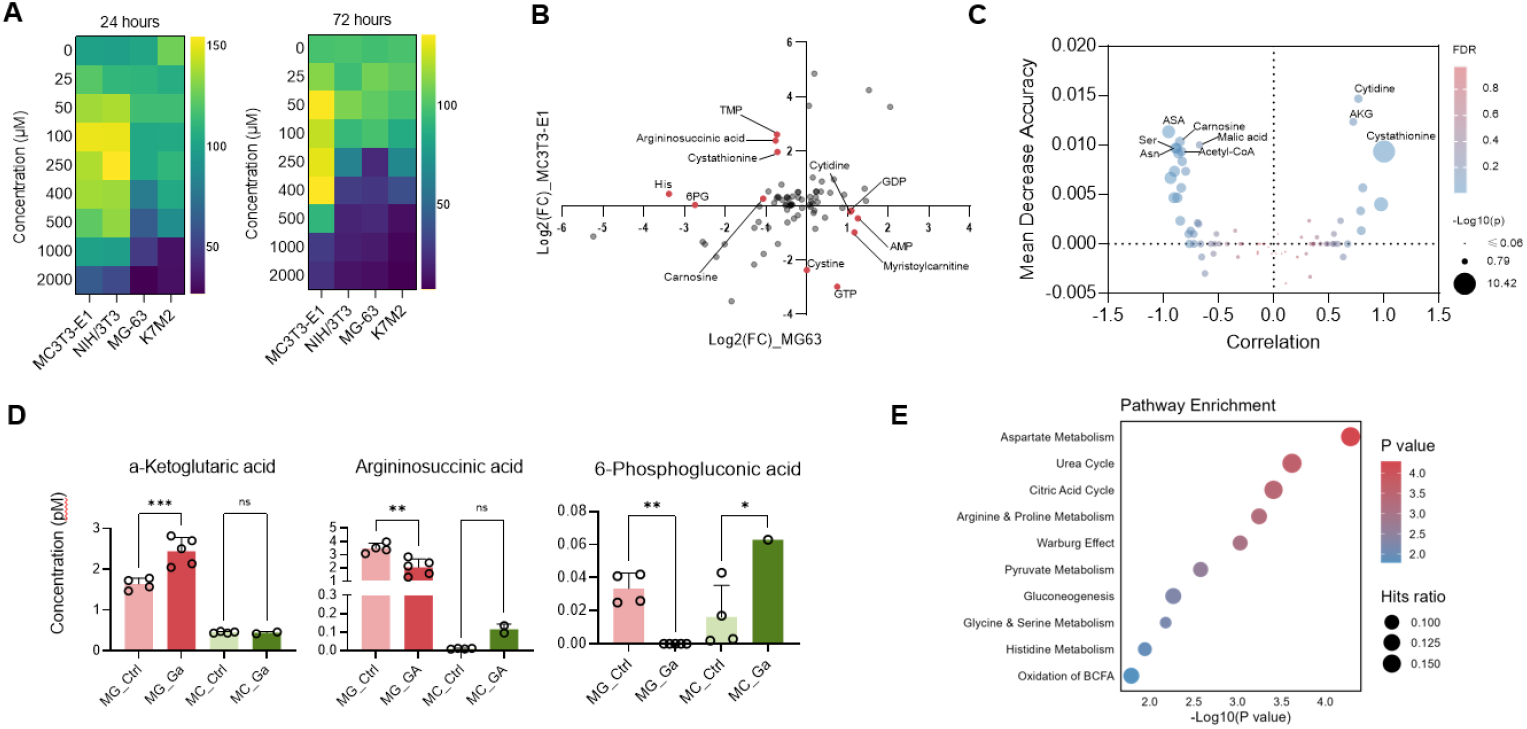
Osteosarcoma and osteoblast exhibit distinct metabolic and survival behaviors under gallium stimulation. **(A**) Cell viability test for MC3T3-E1 pre-osteoblasts, NIH/3T3 fibroblast, MG-63 osteosarcoma cells, and K7M2 osteosarcoma cells treated by gallium nitrate for 24 hours and 72 hours. **(B)** The metabolite profiles of both cells were compared by fold change (FC) and summed together to form a scatter plot. The metabolites with significant (|Log2(FC)| >1) inverse directional changes were labeled. **(C)** Significant different metabolites in-between 2 cell types selected by pattern search (x-axis) and random forest (y-axis) **(D)** Metabolite level of α-ketoglutarate, argininosuccinic acid, and 6-phosphogluconic acid. One-way ANOVA test, **P < 0.01, *P < 0.05. **(E)** Pathway enrichment analysis of the inversely altered metabolites, utilizing the HMDB database. Bubble size represents the enrichment ratio, while the x-axis and color gradient indicate the P-value.

To elucidate the mechanisms driving this selectivity, we performed targeted metabolomic profiling, quantifying 83 metabolites in MG-63 cells and 78 in MC3T3-E1 cells, encompassing amino acids, nucleotides, and organic acids. We plotted the log-transformed fold changes of metabolites in the MG-63 cells against those in the MC3T3-E1 cells to isolate specific compounds that were significantly upregulated in one cell type while concurrently downregulated in the other. The screened metabolites were characterized by their inverse directional changes (Fig. 1B). Specifically, 6-phosphogluconic acid (6-PG), argininosuccinic acid (ASA), carnosine, cystathionine, histamine, and TMP were decreased in sarcoma but increased in osteoblasts, while cytidine, GDP, GTP, AMP, and myristoylcarnitine were increased in sarcoma but decreased in osteoblasts. Subsequent pattern search and Random Forest analyses (Fig. 1C) confirmed that these inversely regulated metabolites, particularly cytidine and α-ketoglutarate (α-KG), serve as the primary discriminators separating the malignant metabolic response from the normal adaptive response.

### Gallium disrupts the cytoskeletal structure and blocks sarcoma cell growth and migration

Given the metabolic crisis induced by gallium, we next investigated its impact on the physical hallmarks of malignancy: migration and invasion. Cell motility was assessed using a wound-healing assay (Fig. 2B and Supplementary Fig. S2A). While the inhibitory effect of gallium was time-dependent— demonstrating a gradual onset over the initial hours—a significant reduction in migration area was established by 15 hours (P < 0.05) and became further pronounced at 24 hours (P < 0.01). This loss of motility was paralleled by a near-total abrogation of invasive capacity. In transwell invasion assays (Fig. 2C), gallium treatment reduced the viability of invading cells by over 90% compared to controls (P < 0.0001), effectively trapping the osteosarcoma cells despite the presence of an FBS chemoattractant in the lower chamber.

**Fig. 2.**
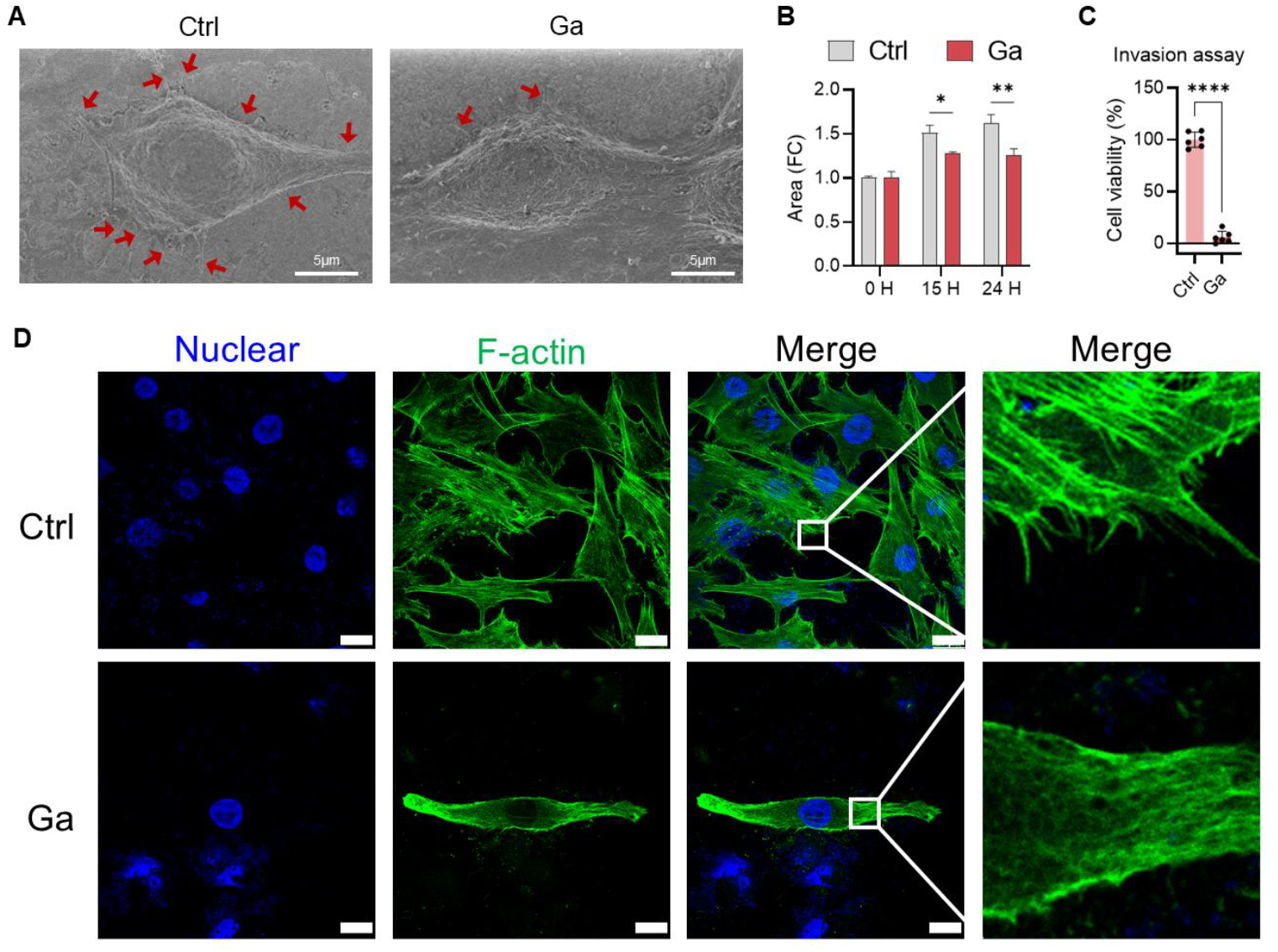
Gallium disrupts the cytoskeletal structure and blocks sarcoma migration. **(A)** SEM image of MG-63 cells, control group and group with gallium for 24 hours. Red arrows point the filopodia. **(B)** Quantified area changes of cell migration assay. One-way ANOVA, **P < 0.01, *P < 0.05. **(C)** Cell viability of the lower chambers from the invasion assay after 96 hours. T-test, ****P < 0.0001. **(D)** Images of actin and nuclei. Scale bar = 2 μm.

, We examined cell surface morphology via scanning electron microscopy (SEM) to elucidate the structural basis underlying this immobility. Control cells exhibited a phenotype typical of highly motile osteosarcoma, characterized by extensive membrane ruffling and numerous filopodial extensions (Fig. 2A, red arrows). In stark contrast, gallium-treated cells displayed a “smooth” phenotype with a marked disappearance of filopodia. Notably, while the cell membrane remained intact (Supplementary Fig. S2C), the cellular machinery required for traction and sensing was dismantled.

Confocal microscopy of F-actin (phalloidin) and nuclei (DAPI) revealed that this morphological loss stems from profound cytoskeletal disorganization (Fig. 2D). Control cells displayed a finely ordered network of F-actin stress fibers and distinct filopodial protrusions. However, gallium treatment induced a chaotic and “blurred” cytoskeletal state. We observed a specific loss or cleavage of actin filaments surrounding the nucleus, accompanied by aberrant actin aggregation at the cell periphery (cortical collapse). Furthermore, nuclear integrity was severely compromised; unlike the intact nuclei of controls, gallium-treated cells exhibited nuclear fragmentation and scattered chromatin, indicative of severe genomic instability or late-stage apoptosis driven by the cytoskeletal collapse.

### Gallium alters energy metabolism and creates accumulation of nucleosides

, We categorized the significantly altered metabolites in MG-63 cells (Fig. 3A) to map the metabolic landscape of gallium-induced toxicity. The resulting profile revealed a distinct metabolic bifurcation: while intermediates of energy production and biosynthesis—specifically within glycolysis and the pentose phosphate pathway (PPP)—were markedly downregulated, there was a simultaneous, paradoxical upregulation of nucleosides and nucleotides. Consistent with an energy crisis model, gallium treatment led to a systemic depletion of glycolytic and PPP intermediates. As modelled in Fig. 3C, we observed significant reductions in upstream metabolites such as glucose-6-phosphate (G6P) and fructose-6-phosphate (F6P), alongside critical downstream effectors like 3-phosphoglycerate (3PG). Notably, the depletion of 6-phosphogluconate (6PG) and ribose-5-phosphate (R5P) confirmed the shutdown of the PPP, effectively severing the supply of NADPH for redox homeostasis and phosphoribosyl pyrophosphate (PRPP) for de novo nucleotide synthesis.

**Fig 3.**
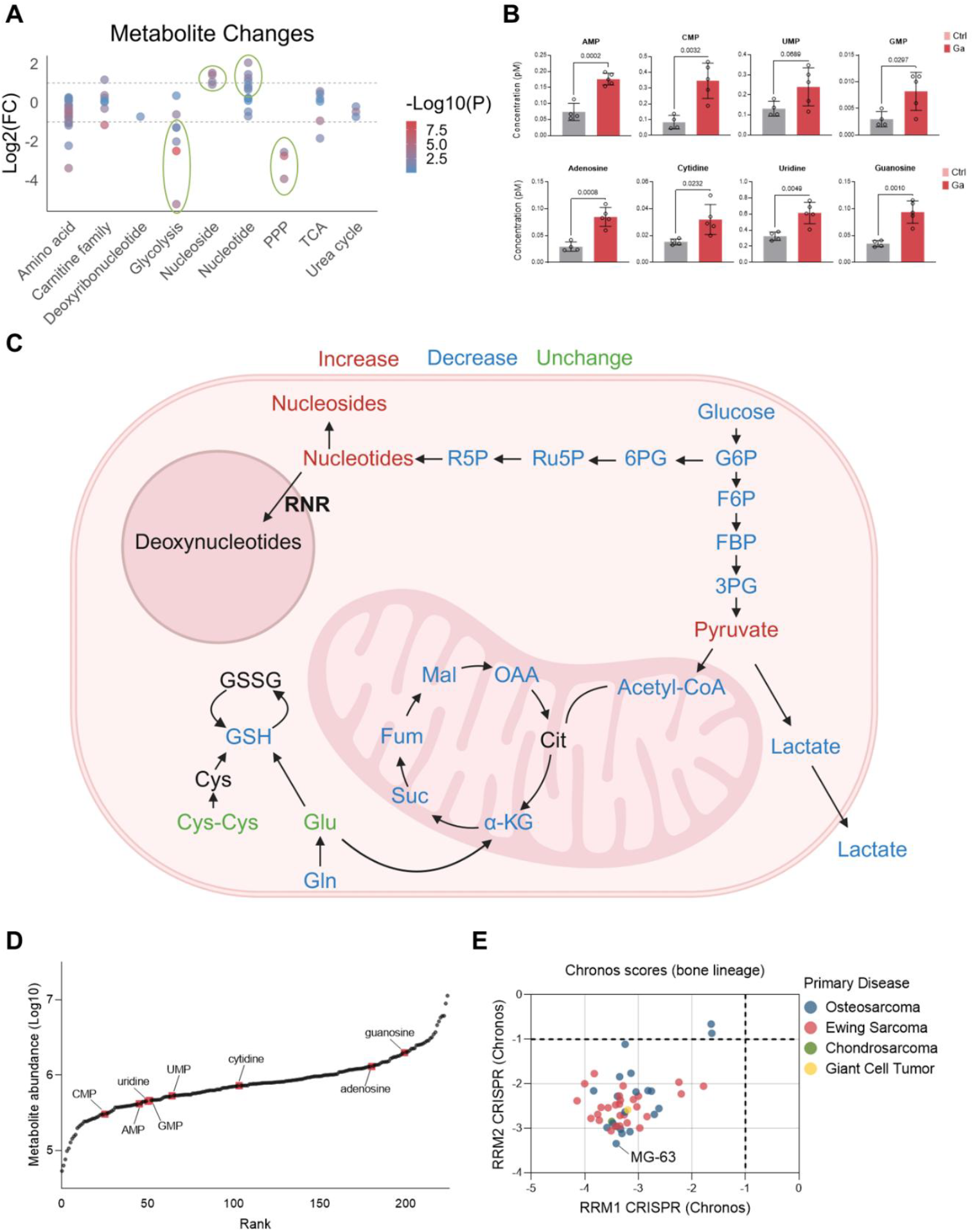
Gallium disrupts central carbon metabolism and induces a ribonucleotide utilization bottleneck. **(A)** Categorical classification of metabolic alterations. Categorical scatter plot of metabolite fold changes in MG-63 cells highlighting the bifurcation between downregulated energy pathways and upregulated nucleotide pools. **(B)** Quantitative profiling of nucleotides and nucleosides showing significant accumulation following Ga treatment. Data: Mean ± SD (n≥4), unpaired t-test. **(C)** Schematic model of the metabolic blockade: Ga suppresses Glycolysis/PPP but causes a backlog at the RNR step, preventing DNA precursor synthesis. **(D)** Waterfall plot of the abundance of metabolites in MG-63 cells, where labeled ribonucleotides and nucleosides reveal their discrete distribution characteristics in an undisturbed state. **(E)** CRISPR-dependent (Chronos) rating scatter plot of bone cancer cell lines on RRM1 and RRM2 subunits.

Despite the suppression of the PPP (the primary synthesis pathway), the intracellular pools of purines and pyrimidines were not depleted; rather, they massively expanded. Quantification of individual species (Fig. 3B) revealed a significant accumulation of monophosphate nucleotides—AMP, CMP, UMP, and GMP—in the gallium-treated cells. The backlog extended to their corresponding nucleosides, with adenosine, cytidine, uridine, and guanosine levels increasing by 2-to 4-fold compared to controls. The accumulation of these substrates, observed under conditions of downregulated biosynthesis, strongly suggests that downstream utilization is impaired rather than synthesis being upregulated. The critical conversion of ribonucleotides into deoxyribonucleotides is catalyzed by RNR.

To contextualize the metabolic vulnerability of MG-63 cells, we examined their basal metabolite landscape (Fig. 3D) using data extracted from the DepMap (https://depmap.org/portal/) [19]. Despite diverse baseline abundances, gallium treatment induced a systemic, paradoxical accumulation of ribonucleotides, which occurred even as upstream precursors like R5P were depleted. This signature identifies a profound metabolic bottleneck, pointing directly to the functional impairment of the RNR-mediated step. We validated this dependency utilizing large-scale CRISPR screening data (Fig. 3E).

MG-63 cells proved highly representative of this specific vulnerability, exhibiting an extreme survival dependency on both the RRM1 and RRM2 subunits of RNR, with Chronos scores significantly below −1. This acute genetic reliance on the RNR-dependent supply of deoxynucleotides provides a compelling biological basis for the gallium-induced metabolic failure. By aligning our metabolomic evidence with the specific genetic sensitivities of MG-63 cells, we established RNR as the primary enzymatic suspect for gallium’s selective cytotoxicity, setting the stage for direct molecular verification.

### Iron but not ferroptosis inhibitors can inhibit cell death

Having identified RNR—an iron-dependent enzyme—as the primary metabolic bottleneck, we hypothesized that gallium exerts its toxicity via “functional iron starvation” rather than primary oxidative damage. To test this, we performed elemental mapping and rescue assays. SEM-EDS analysis confirmed that MG-63 cells robustly internalize gallium (Fig. 4A). Crucially, the addition of exogenous iron (Fe^3+^) did not competitively inhibit gallium uptake. As shown in the merged elemental maps and spectral quantification (Fig. 4B), cells in the Ga + Fe condition displayed strong, co-localized signals for both elements. This indicates that gallium and iron are internalized simultaneously, likely through shared but non-saturating transport mechanisms. Despite the persistent presence of intracellular gallium, iron supplementation provided a complete rescue of cellular viability (Fig. 4C). The addition of equimolar concentrations of iron (250 μM) to gallium-treated cells restored viability to control levels (P < 0.0001). The results suggests that gallium toxicity is not irreversible but rather stems from a competitive equilibrium at specific molecular targets.

**Fig. 4.**
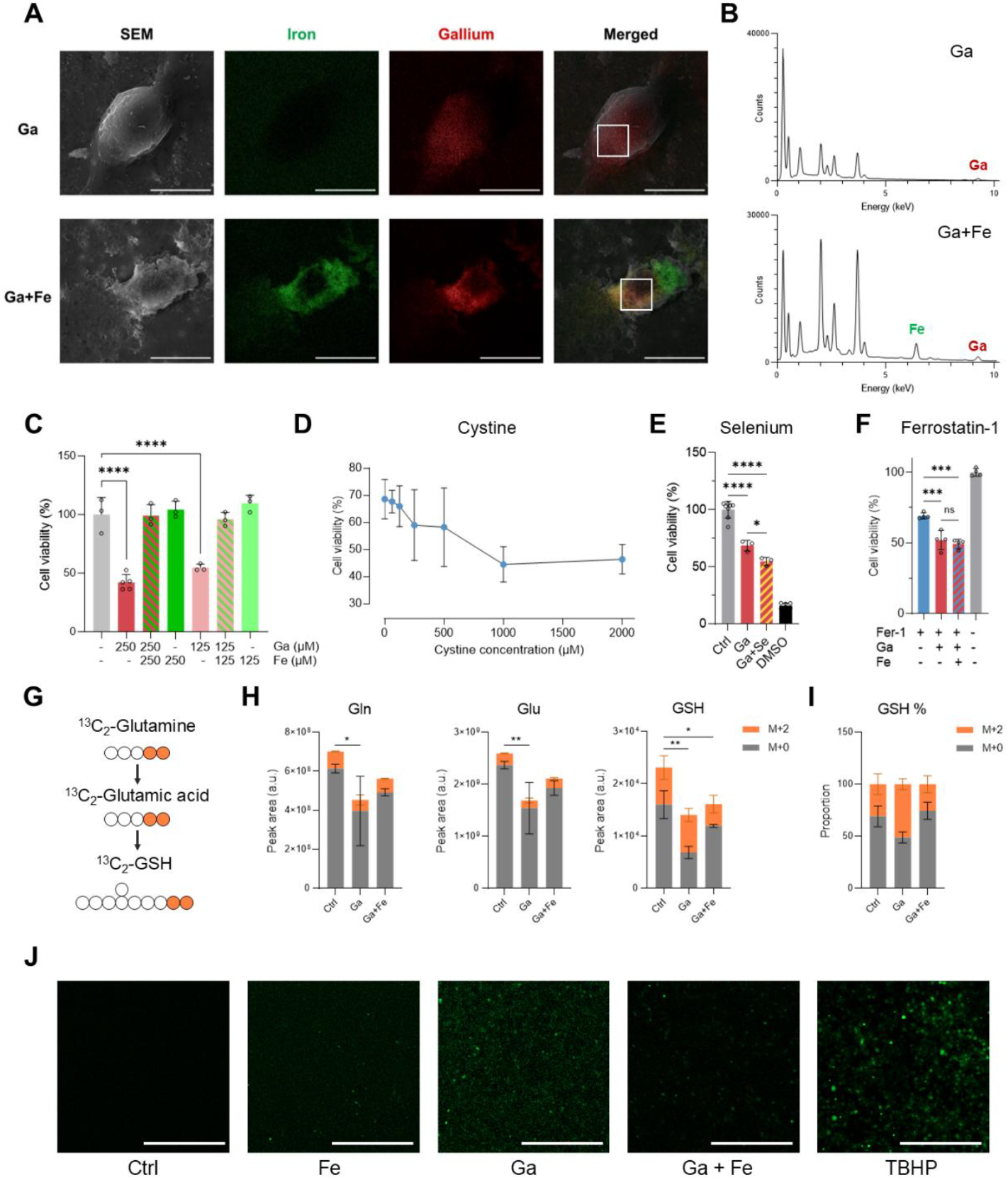
Gallium-induced cytotoxicity is mediated by iron competition rather than canonical ferroptosis. **(A)** Representative Scanning Electron Microscopy (SEM) images and corresponding Energy Dispersive X-ray Spectroscopy (EDS) elemental maps showing the distribution of iron (green) and gallium (red) in MG-63 cells treated with Gallium (Ga) alone or in combination with Iron (Ga + Fe). Scale bars: 10μm. **(B)** EDS spectra quantifying intracellular Ga and Fe content. The profiles confirm robust Ga internalization in both groups, whereas significant Fe signals are restricted to the Ga + Fe supplemented condition. **(C)** Dose-response survival assay of cells treated with Ga (125–250μM) in the presence or absence of equimolar Fe. Fe supplementation significantly reverses Ga-induced lethality. **(D–F)** Viability assays evaluating Ga-induced toxicity under treatment with: **(D)** varying concentrations of cystine; **(E)** selenium; and **(F)** ferrostatin-1. The lack of significant rescue indicates that classical ferroptosis is not the primary mechanism of cell death. **(G)** Isotope flow diagram of ^13^C-glutamine in cells. **(H)** Intracellular glutamine, glutamate, and GSH levels (M+0 and M+2) after 3 hours of isotope treatment. Two-way ANOVA. **(I)** The proportion of GSH in M+0 and M+2 in each group. The gallium-only group exhibited a higher percentage of M+2 GSH. **(J)** Fluorescent imaging of intracellular reactive oxygen species levels under indicated conditions. Scale bar = 100 μm. TBHP serves as a positive control for ROS induction. (Data Analysis) Statistical significance for C, E, and F was determined via one-way ANOVA: ****P<0.0001, ***P<0.001, **P<0.01, *P<0.05.

Given that intracellular iron accumulation often triggers ferroptosis, and considering that gallium mimics iron, we investigated whether gallium induces a pseudo-ferroptosis. Supplementation with cystine, the precursor for glutathione (GSH) synthesis, failed to rescue gallium toxicity and exhibited a slight inhibitory trend at high concentrations (Fig. 4D). Furthermore, selenium, which is an essential cofactor for glutathione peroxidase 4 (GPX4) activity, did not improve cell survival (Fig. 4E). Treatment with ferrostatin-1 (Fer-1), the canonical ferroptosis inhibitor, also showed no significant protective effect against gallium (Fig. 4F).

To further investigate the mechanism by which gallium undermines cellular antioxidant defenses, we performed stable isotope tracing using 13C2-glutamine. The metabolic flow was monitored from 13C2-glutamine (Gln) through 13C2-glutamate (Glu) to the final synthesis of 13C2-glutathione (GSH) (Fig. 4G). Intracellular levels of both Gln and its primary metabolite, Glu, were significantly reduced in the Ga-treated group compared to the control (Fig. 4H). However, the proportion of the M+2 isotopologue remained unchanged for both glutamine and glutamic acid (Supplementary Fig. 4A). Absolute GSH levels (both M+0 and M+2 fractions) decreased markedly upon gallium treatment and gallium-iron combination treatment, with the gallium-only treatment showing the most significant decrease (Fig. 4H). As GSH is crucial for redox homeostasis, we independently verified the mass spectrometry data using a biochemical GSH assay kit. Consistent with previous experiments, the results confirmed that GSH levels decreased due to gallium exposure [20,21].

Interestingly, while total GSH plummeted, the proportion of ^13^C_2_-labeled GSH (M+2) was significantly higher in the Ga-only group compared to other groups (Fig. 4I). This dynamic indicates that while the cell compensatory shunts remaining glutamine toward GSH synthesis to combat stress, the overall metabolic flux remains insufficient to maintain homeostatic levels. Although the addition of iron also reduced overall GSH content, it did not result in an acceleration of the synthesis pathway from glutamate to GSH. Finally, to visually assess ROS generation, an intracellular ROS assay was performed utilizing fluorescence microscopy (Fig. 4J). Gallium substantially increased the intensity of intracellular ROS, whereas the addition of iron ions mitigated this ROS accumulation. Tert-butyl hydroperoxide (TBHP) was applied as a positive control to induce ROS.

Connecting our prior observation of ribonucleotide accumulation to our screening of iron-dependent enzymes, we identified RNR as the critical enzymatic bottleneck responsible for the downstream processing of these metabolites (Fig. 3C). Building upon the discovery that gallium induces a functional iron deficiency at the RNR active site, we next examined the downstream consequences on genomic stability. Alkaline comet assays revealed that Ga treatment triggers extensive DNA fragmentation in MG-63 cells (Fig. 5A). While control nuclei remained intact, gallium-treated cells exhibited prominent comet tails, signifying a marked accumulation of single- and double-strand DNA breaks. This structural damage is highly consistent with a substrate starvation model: by inhibiting RNR, gallium depletes the deoxyribonucleotide (dNTP) pools strictly required by high-fidelity DNA polymerases to execute routine replication and repair.

**Fig 5.**
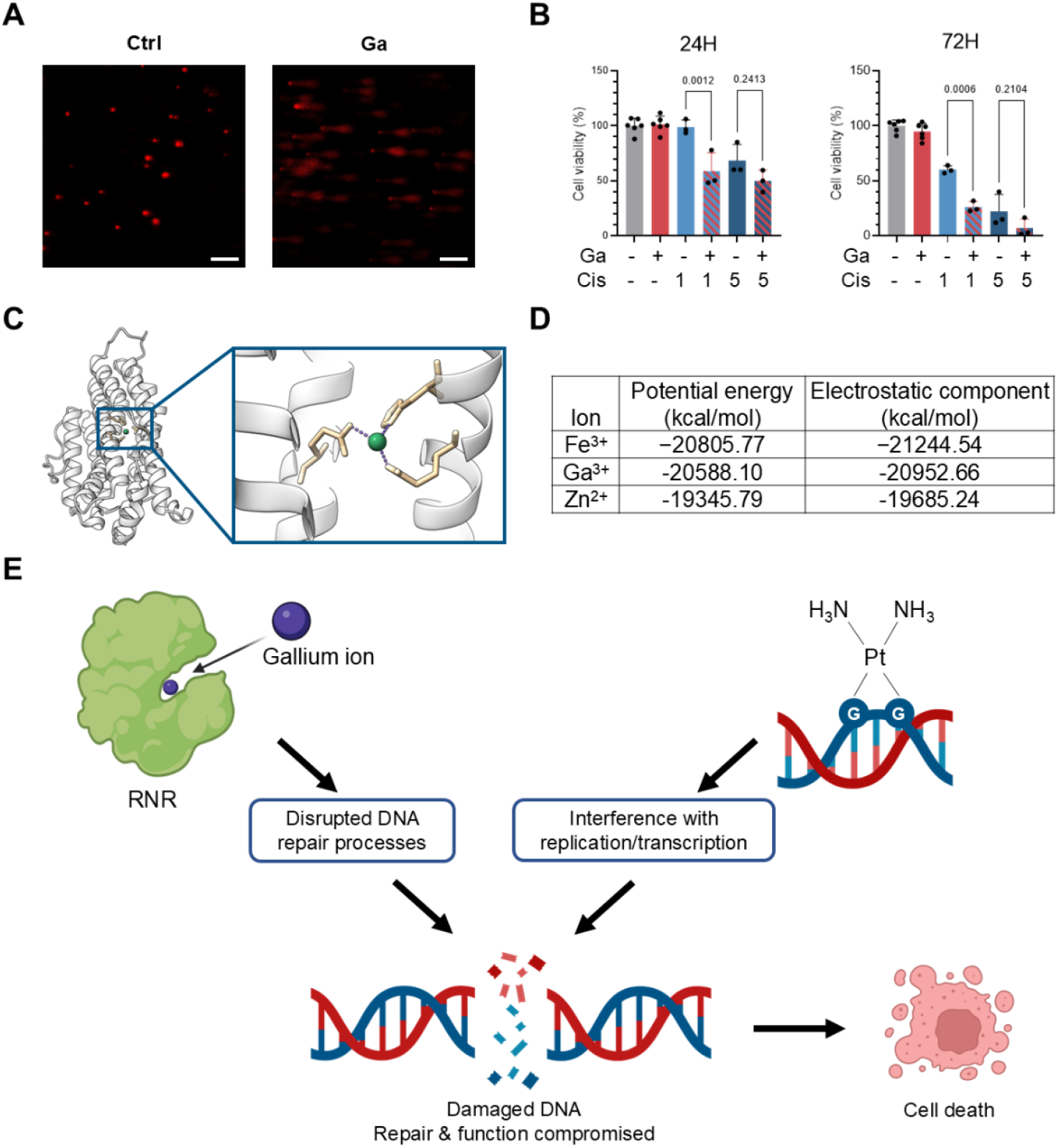
Gallium disrupts DNA repair by binding to RNR and synergizes with cisplatin. **(A)** Representative images from an alkaline comet assay in MG-63 cells. Scale bar = 5 μm. **(B)** Cell viability analysis of MG-63 cells treated with gallium (Ga) at 100 μM and cisplatin (Cis) at indicated concentrations (1 μM and 5 μM), or a combination of both for 24 and 72 hours. Data are presented as mean ± SD; p-values indicate comparisons between Cis-only and Ga+Cis cohorts, one-way ANOVA test. **(C)** Molecular visualization of the metal center within the human RNR subunit M2 (PDB: 2UW2). The inset highlights the coordination environment of the metal ion (green sphere) by key amino acid residues. Figure was generated by ChimeraX [22]. **(D)** A table presenting the calculated potential energy and electrostatic components for Fe^3+^, Ga^3+^, and Zn^2+^ ions substituted into the RNR active site. The more negative potential energy compared to the native suggests a thermodynamically favorable, stable binding that would result in enzyme inactivation. **(E)** Schematic illustrating the cooperative action of gallium and cisplatin targeting DNA repair pathways.

To therapeutically exploit this metabolic vulnerability, we evaluated the efficacy of gallium in combination with the DNA-damaging agent cisplatin (Cis). Cell viability assays demonstrated a significant synergistic effect. At 72 hours, the addition of gallium to a low-dose cisplatin regimen (1 μM) resulted in a substantially more pronounced reduction in cell survival compared to cisplatin monotherapy. This synergy indicates that gallium functions as a metabolic DNA repair inhibitor; by throttling the supply of dNTPs through RNR blockade, gallium effectively prevents the cell from synthesizing the DNA patches required to repair cisplatin-induced platinum adducts.

To elucidate the molecular basis of this gallium-induced RNR inhibition, we performed in silico energy minimization on the human RNR active site (PDB: 2UW2) (Fig. 5C). Native Fe^3+^ established a stability benchmark with a total potential energy (E_total_) of −20,805.77 kcal/mol and a dominant electrostatic component of −21,244.54 kcal/mol (Fig. 5D). Substitution with Ga^3+^ revealed remarkable structural mimicry. The Ga^3+^-RNR complex achieved a converged E_total_ of −20,588.10 kcal/mol, retaining 98.9% of the native electrostatic stability (−20,952.66 kcal/mol). In contrast, Zn^2+^ exhibited significantly lower affinity, with E_total_ dropping to −19,345.79 kcal/mol. The substantial energy gap stems from the lower formal charge and geometric mismatch of Zn^2+^ within the octahedral pocket. These findings demonstrate that gallium can act as a potent structural decoy, outcompeting native iron through thermodynamic mimicry to inactivate RNR, whereas zinc fails to maintain sufficient interaction density to effectively disrupt the enzymatic center.

Ultimately, the combination of gallium and cisplatin demonstrates significant clinical potential by simultaneously targeting DNA integrity and the metabolic supply chain required for genomic repair (Fig. 5E). Specifically, gallium’s inhibition of RNR creates a dNTP substrate starvation that prevents the repair of cisplatin-induced DNA lesions, thereby overwhelming the cancer cell’s innate survival mechanisms.

## Discussion

Over the past decades, gallium has attracted sustained interest in both academic and clinical fields due to its broad anti-cancer, anti-inflammatory, and anti-bacterial activities. Clinically, gallium salts and gallium-based compounds have been explored for the treatment of malignancies and bone-related disorders, capitalizing on gallium’s ability to mimic ferric iron while remaining redox-inactive. [23–25]. More recently, gallium has also been incorporated into biomaterials to endow implants with anti-tumor and anti-microbial properties, particularly in the context of bone repair [1,2]. Despite this growing translational interest, the molecular mechanisms underlying gallium cytotoxicity remain incompletely defined, limiting the rational optimization of gallium-based therapeutics. Previous studies have largely attributed gallium toxicity to broad concepts like iron competition or iron deprivation [26]. However, these descriptions are mechanistically indirect, failing to explain why gallium exhibits selective toxicity toward highly proliferative cancer cells while sparing normal tissues, or how this metabolic stress translates into cell death.

To address this gap and identify new strategies to break cancer drug resistance, we employed an interdisciplinary approach centered on metabolomics. This strategy revealed a defining metabolic signature of gallium treatment in sarcoma cells: a profound bifurcation of central carbon metabolism. We observed a systemic depletion of intermediates in glycolysis and the pentose phosphate pathway (PPP), including the critical depletion of ribose-5-phosphate (R5P), which starves the cell of the 5-carbon sugars required for de novo nucleotide synthesis. Crucially, despite this upstream synthetic shutdown, our metabolomic profiling uncovered a novel accumulation of ribonucleotide monophosphates (AMP, CMP, UMP, GMP) and nucleosides. The paradoxical accumulation identifies a profound downstream utilization blockade, providing an unusually high-resolution metabolic readout that directly infers enzyme-level dysfunction at ribonucleotide reductase (RNR). This observation offers an unusually high-resolution metabolic readout, allowing inference of enzyme-level dysfunction directly from intracellular metabolite pools rather than indirect proliferation or DNA synthesis markers [27,28].

RNR is uniquely vulnerable to metal perturbation due to its absolute requirement for a correctly coordinated di-iron center to sustain radical-based catalysis. Our computational modeling demonstrates that Ga3+ exhibits near-perfect thermodynamic and electrostatic mimicry of Fe^3+^ within the RNR metal-binding pocket, achieving 98.9% of native binding stability. This high-fidelity structural decoy enables gallium to outcompete iron for enzyme occupancy while remaining redox-inactive, irreversibly disabling RNR function. We utilized SEM-EDS elemental mapping to visually validate this interaction, demonstrating that iron and gallium are internalized simultaneously and co-localize within the cell. The dynamic relationship between the cellular uptake and retention of these ions helps explain the precise cellular response to gallium. Importantly, exogenous iron supplementation rescued cell viability despite continued gallium uptake (Fig. 4C), confirming that gallium toxicity operates through a reversible competitive equilibrium at functional sites like RNR rather than through nonspecific metal poisoning. We therefore define gallium-induced cytotoxicity as a state of “functional iron starvation,” wherein iron is present intracellularly but rendered biochemically inaccessible to essential enzymes. The dynamic relationship between cellular uptake and retention of iron and gallium ions was demonstrated in a recently published study [29], which, along with our experimental results, can help explain cellular responses to gallium. While some studies have observed synergistic cytotoxicity with gallium and attributed the resulting cell death to elevated ROS and ferroptosis [10], our data suggest this may be an incomplete conclusion. The isotope tracing and rescue assays indicate that gallium-induced ROS accumulation and DNA repair inhibition are more likely independent consequences of a broader metabolic collapse. The functional iron starvation imposed by gallium directly halts the dNTP supply line, leading to the DNA fragmentation observed in our comet assays, independent of classical ferroptotic pathways.

The metabolic blockade imposes a precise and exploitable opportunity to address an unmet clinical need in oncology: overcoming chemoresistance. By inhibiting RNR, gallium restricts de novo dNTP synthesis, effectively functioning as a metabolic DNA repair inhibitor. When combined with the DNA-damaging agent cisplatin, gallium prevents cancer cells from generating the nucleotide substrates required to repair platinum-induced DNA adducts, converting otherwise repairable lesions into irreversible genomic damage. The dual-hit strategy—simultaneous induction of DNA damage and suppression of repair capacity—provides a robust mechanistic basis for breaking platinum resistance. As this approach selectively targets iron-dependent nucleotide metabolism, it preferentially sensitizes highly proliferative osteosarcoma cells while sparing normal osteoblasts, establishing a favourable therapeutic window in bone tissue.

Beyond its immediate therapeutic relevance, the presented study highlights RNR as a central metabolic checkpoint linking iron homeostasis to DNA repair and cell fate. More broadly, it establishes the interdisciplinary application of metabolomics to restrict nucleotide supply as a rational strategy to enhance the efficacy of genotoxic chemotherapy. It is worth noting that, due to metabolic differences, the dosage of gallium in iron-deficient cancer patients will need to be controlled with extreme caution. In conclusion, by combination of metabolomics and SEM-EDS elemental mapping, we identified RNR as the primary molecular target of gallium. These insights establish gallium as a potent DNA repair inhibitor, demonstrating significant clinical potential in combination with cisplatin to overwhelm sarcoma cell survival mechanisms and overcome therapeutic resistance.

## Materials and Methods

### Cell culture studies

MG-63 (CRL-1427), MC3T3-E1 (CRL-2593, subclone 4), K7M2 (CRL-2836), and NIH/3T3 (CRL-1658, ECACC 93061524) cell lines were cultured and studied. MG-63, K7M2, and MC3T3-E1 cell lines were used for fewer than 7 passages, NIH/3T3 was used for fewer than 16 passages. RPMI-1640 and MEMα were applied in this study because neither of these two culture media contains added iron salts, which may be a huge factor affecting experimental results. To maintain the consistent nutritional conditions for metabolomics experiments, cells were cultured in the same culture media. To Meet MC3T3-E1 growth conditions, in cell viability assay, GSH assay, and metabolomics part, cells were cultured with Minimum Essential Medium Alpha (MEMα) (Thermo Fisher) supplemented with 10% (vol/vol) Fetal Bovine Serum (FBS, Sigma-Aldrich), 1% (vol/vol), and Penicillin Streptomycin (P/S, Gibco) inside a 5% CO_2_ incubator at 37 °C. RPMI-1640 (Thermo Fisher) with 10% (vol/vol) FBS and 1% (vol/vol) P/S was used as media in fluorescent staining, iron ion intervention, invasion assay, and migration assay with MG-63 cells.

### Cell viability assay

Cell viability was assessed by Cell Counting Kit-8 (Dojindo) assay. Cells were seeded at 4000 cells/well in 96-well plates with 100 μl culture medium. After 24 hours, another 100 μl culture media with a gradient concentration of gallium nitrate (Sigma-Aldrich, ≥99.9%) was added. The final concentrations of gallium ions were 0, 25, 50, 100, 250, 400, 500, 1000, and 2000 μM, respectively. 5% DMSO was used as positive control. The assay was conducted according to the manufacturer’s instructions after 24 hours and 72 hours adding the gallium contained media.

### Fluorescent staining of actin and nucleus

MG-63 cells were seeded in chamber slide (Slide 8 Well, Ibidi) overnight to attach. Media was replaced with the gallium-contained media (250 μM). Media without gallium was used for control group. After 24 hours treating, cells were washed with PBS and then fixed for 10 min at room temperature with 4% paraformaldehyde in PBS. The fixed cells were permeabilized for 20 min at room temperature with permeabilization buffer (0.1% Triton X-100 in PBS), followed by two washes with PBS. The cells were incubated at room temperature in the dark for 20 min with phalloidin (Alexa Fluor 488, Thermo Fisher) and DAPI (DAPI solution, Thermo Fisher) according to the manufacture’s protocol. All fluorescence images were captured using a confocal microscope (STELLARIS SP8, Leica). Nuclei were excited at 405nm, emission spectrum was collected in range from 425-507nm. Actin was excited at 497nm, emission spectrum was collected in range from 507-650nm.

### Cell invasion and migration assay

In the invasion assay, 6-well plates with inserts were employed. The inserts were coated with filter-sterilized gelatin derived from porcine skin (Sigma) to simulate the extracellular matrix. MG-63 cells were seeded at a density of 6 × 10^5^ cells per well onto the gelatin-coated inserts and allowed to attach overnight. The lower chamber of the plate was filled with complete media. Once cell attachment was established, the media in the upper chamber (the insert) was replaced with media lacking FBS. In the experimental groups, the media used (both upper and lower chambers) contained gallium at a concentration of 125 μM, while the control group received media without gallium. To quantify the migrated cells, a cell viability assay using CCK-8 was performed in the lower chamber after 96 hours.

In the migration assay, 12-well plates were employed. MG-63 cells were seeded at a density of 1.2 × 10^5^ cells per well. Cells were allowed to attach overnight and, then the media of experimental groups was changed into the media with gallium ion (250 μM). Media in control groups was changed into fresh media. After 24 hours, cells were scratched by 200 μL pipette tips. Cells were washed with PBS for 3 times and changed into new media (with or without gallium). Images were taken by microscope (C-P20, OPTIKA) at 0, 15, and 24 hours after the scratching.

### Intracellular metabolites collection and preparation

Both MG-63 and MC3T3-E1 cells were seeded in 24-well plates, 0.8 × 10^5^ cells per well with 500 μL culture media. After 24 hours, media with gallium was added (500ul) and the final concentration of Ga^3+^ was 400 μM. Cells were cultured for another 24 hours. Before the extraction, cell culture media was removed, and cells were washed with ammonium bicarbonate (NH_4_HCO_3_, Sigma-Aldrich, ≥99.5%) solution (75mM, 37°C). Cells were then quenched in cold 80% v/v methanol (CH_3_OH, Sigma-Aldrich, ≥99.9%) and harvested by scraping. The harvested samples were centrifuged at 1000 RCF at room temperature for 5 minutes, and the supernatant was collected. The collected supernatant was dried by vacuum centrifuge. Afterward, 10 μL of the isotopically labeled internal standard mix was added to dried samples, followed by 90 μL of methanol. Reconstituted samples were transferred to glass vials and applied for liquid chromatography-mass spectrometry (LC-MS) analysis.

### LC-MS based metabolomics

A Vanquish Core HPLC system (Thermo Scientific) coupled with an Orbitrap Exploris 120 (Thermo Scientific) mass spectrometer was used for the LC-MS analysis. Two different chromatography methods were used. Two chromatographic methods were employed depending on the ionization mode of mass spectrometer. The chromatographic separation was carried out on an Atlantis Premier BEH Z-HILIC Column, 1.7 µm, 2.1 mm X 50 mm (Waters). For the postive ionization mode the column was maintained at 40 °C, and the sample injection volume was 2 µL. The mobile phase consisted of phase A - 0.15% formic acid (v/v) in water and phase B - 0.15% formic acid (v/v) in 85% acetonitrile (v/v) with 10 mM ammonium formate. The gradient elution with a flow rate of 0.4 mL/min was performed for a total analysis time of 17 min. For the negative ionization mode the column was maintained at 40 °C, and the sample injection volume was 2 µL. The mobile phase consisted of phase A - 15 mM Ammonium bicarbonate, pH 9.00 in water and phase B - 15 mM Ammonium bicarbonate, pH 9.00 in 90% acetonitrile (v/v). The gradient elution with a flow rate of 0.4 mL/min was performed for a total analysis time of 15 min.

The Orbitrap Q Exploris 120 (Thermo Scientific) mass spectrometer was operated in either positive or negative electrospray ionization mode, spray voltage 3.5 kV, aux gas heater temperature 400 °C, capillary temperature 350 °C, aux gas flow rate 12, sheath gas flow rate 50. The metabolites of interest were analyzed using a full MS scan mode, scan range m/z 50 to 600, resolution 30000. The Trace Finder 4.1 software (Thermo Scientific) was used for data processing. A seven-point linear calibration curve with internal standardization and 1/x weighing was constructed for the quantification of the metabolites.

### Bioinformatics and Dependency Analysis (DepMap)

Basal Metabolomics Waterfall Plot To characterize the baseline metabolic landscape of osteosarcoma, we utilized the Metabolomics (Public 25Q3) dataset via the DepMap Data Explorer 2.0 (https://depmap.org/portal). A Waterfall plot was generated specifically for the MG-63 cell line (Model ID: ACH-000359). Metabolite abundance was represented on the Y-axis as -transformed intensities, with individual metabolites ranked along the X-axis by their basal abundance. Genetic Dependency Analysis (Chronos Scores) The genetic dependency of bone cancer lineages on the Ribonucleotide Reductase (RNR) complex was assessed using CRISPR (DepMap Public 25Q3+Score, Chronos) data. We extracted the Chronos scores—a measure of gene essentiality where lower scores indicate higher dependency—for the RRM1 and RRM2 subunits. A scatter plot was constructed to visualize the co-dependency of these subunits across the Bone lineage. To evaluate the representativeness of our model, the MG-63 cell line was specifically highlighted within the distribution of other bone-derived malignancies. A threshold of −1 was utilized to define common essentiality.

### GSH assay

MG-63 cells were seeded at 8000 cells/well in 96-well plates with 100 μl culture medium. After 24 hours, another 100 μl culture media with gallium nitrate was added to reach the final Ga^3+^ concentration at 400 μM. After a further 24-hour incubation, cellular GSH was tested with a GSH assay kit (Merck, USA) via enzymatic cycling method according to the manufacturer’s instructions. The cells were washed with PBS, followed by the addition of 200 µL Sulfosalicylic acid 5% to avoid oxidation of GSH content. Immediately afterward, the cells were lysed by placing the plate in a sonicator bath (with ice) (Cole-Parmer, USA) for 5 min. Then, all the wells were scraped and collected in a tube followed by centrifuging at 6000 rpm for 10 min. Next, 20 µL of the supernatants were collected and diluted 3 times with GSH assay buffer. Next, 20 µL of each sample was transferred to a transparent 96-well plate and mixed with a reaction mix to start the enzymatic reaction. OD was recorded using the plate reader at 450 nm at two different time points (20 min and 40 min). The results were reported as % of the control.

### Stable isotope tracing with ^13^C_2_-glutamine

MG63 osteosarcoma cells were seeded in 12-well plates at a density of 1.0 × 10^5^ cells per well and cultured in RPMI-1640 medium. Cells were allowed to adhere overnight before treatment. The following day, cells were treated with gallium nitrate (Ga; 250 μM), gallium nitrate in combination with iron(III) nitrate (Ga–Fe; each at 250 μM), or vehicle control. Gallium nitrate (≥99.9%) and iron(III) nitrate (≥99.95%) were purchased from Sigma-Aldrich. After 24 h of treatment, the culture medium was replaced with isotope-labeled medium consisting of RPMI-1640 supplemented with 10% fetal bovine serum and extra 10% ^13^C_2_-glutamine (Cambridge Isotope Laboratories). Cells were incubated in the labeled medium for 3 h to allow metabolic incorporation of the tracer. Metabolism was then quenched by rapidly washing cells with ammonium bicarbonate with a pH of 7, followed by extraction with 80% (v/v) ice-cold methanol. Cell extracts were collected and stored at −80 °C prior to mass spectrometry-based metabolomic analysis. Each experimental condition was performed in triplicate.

### DNA damage assay

The Comet Assay is performed by embedding cells in low-melting-point agarose on a microscope slide, followed by cell lysis in a high-salt, alkaline buffer to remove membranes and proteins, leaving supercoiled DNA attached to the nuclear matrix. The slides are then placed in an alkaline electrophoresis solution to allow DNA unwinding, and an electric field is applied, causing damaged DNA fragments to migrate toward the anode. After electrophoresis, the DNA is neutralized, stained with ethidium bromide (15585011, Thermo Scientific, Massachusetts, USA), and visualized under a fluorescence microscope.

### Intervention of iron ions

To modify the complete media, gallium nitrate and/or iron (III) nitrate (Sigma-Aldrich, ≥99.95%) were added and filter sterilized. The final concentration of gallium ions and/or iron ions used was 125 μM and 250 μM, respectively. MG-63 cells were seeded at a density of 8000 cells per well in 96-well plates with an initial volume of 100 μl of the original culture medium. After 24 hours, the media was aspirated and replaced with media supplemented with the respective/mixed metal ions. Cell viability was assessed using the CCK-8 assay after a further 24-hour incubation period.

### Intracellular ROS evaluation

Intracellular ROS levels were assessed using the DCFDA/H2DCFDA Cellular ROS Assay Kit (Abcam) according to the manufacturer’s protocol. MG-63 osteosarcoma cells were seeded 96-well plates (1 × 10^4^ cells/well) and allowed to attach overnight. Cells were washed with PBS and incubated with 20 µM DCFDA for 40 min at 37°C in the dark, followed by gentle washing with assay buffer. Fluorescent dye– loaded cells were then treated with gallium nitrate (400 µM), iron(III) nitrate (400 µM), their combination (400 µM each), or TBHP (110 µM; positive control), while untreated cells served as a negative control; all treatments were prepared in supplied buffer. After 4 hours of incubation at 37°C, fluorescence was visualized using an inverted fluorescence microscope (LSM 900, Zeiss).

### In Silico Metal Substitution and Energy Minimization

The crystal structure of human RNR was retrieved from the Protein Data Bank (PDB ID: 2UW2). All solvent molecules were removed prior to simulation. The native Fe^3+^ ion at the active site was replaced by Ga^3+^ and Zn^2+^ through direct modification of atomic properties in UCSF ChimeraX, including reassignment of atomic number and atom type, while preserving the original coordination geometry [22]. Hydrogen atoms were added to the structure. Force field parameters and partial charges for the protein were assigned using the AMBER ff14SB force field. Energy minimization was performed using 2,000 steps of conjugate gradient minimization to relieve steric clashes and allow local relaxation of coordinating residues. Total potential energy and its electrostatic and van der Waals components were extracted before and after minimization.

### Statistical analysis

All cell experiments were performed at least triplicate. The data are expressed as mean ± SD. The metabolite profiles were analyzed by MetaboAnalyst 5.0 and GraphPad Prism 9. Metabolomics data was normalized by log transformation (base 10) and auto scaling (mean-centered and divided by the standard deviation of each variable) in MetaboAnalyst 5.0 [30]. One-way ANOVA test was applied with multiple comparisons, compared the mean of each group with the mean of every other groups. Differences were analyzed by two-sided t test. Pathway analysis and enrichment analysis used The Small Molecule Pathway Database as the pathway library. Visualization of the enrichment network of metabolites correlation was performed by Metscape 2 in Cytoscape 3.9.1.

To isolate metabolites that exhibit cell-type-dependent responses to Gallium, we developed a fold-change-based classification workflow (Fig. 1C). First, raw metabolite intensities were normalized. For each of the four biological replicates (n=4) in the Gallium-treated groups (MG-63 and MC3T3-E1), a Fold change value was calculated relative to the average intensity of the corresponding untreated control group. This transformation generated a normalized dataset consisting of 8 data points per metabolite (4 fold change values for MG-63 and 4 fold change values for MC3T3-E1). This dataset served as the input for two parallel analyses, Random Forest classification to distinguish between the two cell types based on their metabolic fold change profiles and pattern search (correlation analysis) to identify metabolites with high importance and strong inverse correlation.

## Ethics approval and consent to participate

All methods were performed in accordance with the relevant guidelines and regulations. No approval of an ethics committee or informed consent was required for this study. No identifiable images from human research participants have been used.

## Supplementary information

Supplementary Material is available.

## Contributions

Jingzhi Fan: Conceptualization, methodology, data curation, validation, visualization, writing—original draft. Annija Vaska: Formal analysis, methodology. Xiping Jiang: Visualization, software. Kristaps Klavins: Funding acquisition, resources, methodology, writing—review and editing, conceptualization, supervision.

## References

1. Kurtuldu F, Mutlu N, Boccaccini AR, Galusek D. Gallium containing bioactive materials: A review of anticancer, antibacterial, and osteogenic properties. Bioact Mater. 2022;17:125–46. 10.1016/j.bioactmat.2021.12.034

2. Mosina M, Kovrlija I, Stipniece L, Locs J. Gallium containing calcium phosphates: Potential antibacterial agents or fictitious truth. Acta Biomater. 2022;150:48–57. 10.1016/j.actbio.2022.07.063

3. Minandri F, Bonchi C, Frangipani E, Imperi F, Visca P. Promises and failures of gallium as an antibacterial agent. Future Microbiol. 2014;9:379–97. 10.2217/fmb.14.3

4. Peng XX, Gao S, Zhang JL. Gallium (III) Complexes in Cancer Chemotherapy. Eur J Inorg Chem. 2022;2022:1–9. 10.1002/ejic.202100953

5. Kubista B, Schoefl T, Mayr L, Van Schoonhoven S, Heffeter P, Windhager R, et al. Distinct activity of the bone-targeted gallium compound KP46 against osteosarcoma cells - Synergism with autophagy inhibition. Journal of Experimental and Clinical Cancer Research. Journal of Experimental & Clinical Cancer Research; 2017;36:1–13. 10.1186/s13046-017-0527-z

6. Timerbaev AR. Advances in developing tris(8-quinolinolato)gallium(iii) as an anticancer drug: Critical appraisal and prospects. Metallomics. 2009;1:193–8. 10.1039/b902861g

7. Chitambar CR. Gallium nitrate for the treatment of non-Hodgkin’s lymphoma. Expert Opin Investig Drugs. 2004;13:531–41. 10.1517/13543784.13.5.531

8. Peng XX, Zhang H, Zhang R, Li ZH, Yang ZS, Zhang J, et al. Gallium Triggers Ferroptosis through a Synergistic Mechanism. Angewandte Chemie - International Edition. 2023;515031. 10.1002/anie.202307838

9. Chen CH, Huang YM, Grillet L, Hsieh YC, Yang YW, Lo KY. Gallium maltolate shows synergism with cisplatin and activates nucleolar stress and ferroptosis in human breast carcinoma cells. Cellular Oncology. Springer Science and Business Media B.V.; 2023;46:1127–42. 10.1007/s13402-023-00804-x

10. Li W, Yang C, Cheng Z, Wu Y, Zhou S, Qi X, et al. Gallium complex K6 inhibits colorectal cancer by increasing ROS levels to induce DNA damage and enhance phosphatase and tensin homolog activity. MedComm (Beijing). John Wiley and Sons Inc; 2024;5. 10.1002/mco2.665

11. Qi J, Qian K, Tian L, Cheng Z, Wang Y. Gallium(iii)-2-benzoylpyridine-thiosemicarbazone complexes promote apoptosis through Ca2+ signaling and ROS-mediated mitochondrial pathways. New Journal of Chemistry. 2018;42:10226–33. 10.1039/c8nj00697k

12. Chang KL, Liao WT, Yu CL, Lan CCE, Chang LW, Yu HS. Effects of gallium on immune stimulation and apoptosis induction in human peripheral blood mononuclear cells. Toxicol Appl Pharmacol. 2003;193:209–17. 10.1016/j.taap.2003.07.004

13. Gallego B, Kaluderovic MR, Kommera H, Paschke R, Hey-Hawkins E, Remmerbach TW, et al. Cytotoxicity, apoptosis and study of the DNA-binding properties of bi- and tetranuclear gallium(III) complexes with heterocyclic thiolato ligands. Invest New Drugs. 2011;29:932–44. 10.1007/s10637-010-9449-8

14. Valiahdi SM, Heffeter P, Jakupec MA, Marculescu R, Berger W, Rappersberger K, et al. The gallium complex KP46 exerts strong activity against primary explanted melanoma cells and induces apoptosis in melanoma cell lines. Melanoma Res. 2009;19:283–93. 10.1097/CMR.0b013e32832b272d

15. Goss CH, Kaneko Y, Khuu L, Anderson GD, Ravishankar S, Aitken ML, et al. Gallium disrupts bacterial iron metabolism and has therapeutic effects in mice and humans with lung infections. Sci Transl Med. 2018;10:1–12. 10.1126/scitranslmed.aat7520

16. Chitambar CR. Gallium and its competing roles with iron in biological systems. Biochim Biophys Acta Mol Cell Res. Elsevier B.V.; 2016;1863:2044–53. 10.1016/j.bbamcr.2016.04.027

17. Yang M, Chitambar CR. Role of oxidative stress in the induction of metallothionein-2A and heme oxygenase-1 gene expression by the antineoplastic agent gallium nitrate in human lymphoma cells. Free Radic Biol Med. 2008;45:763–72. 10.1016/j.freeradbiomed.2008.05.031

18. Chitambar CR. Medical applications and toxicities of gallium compounds. Int J Environ Res Public Health. 2010;7:2337–61. 10.3390/ijerph7052337

19. Arafeh R, Shibue T, Dempster JM, Hahn WC, Vazquez F. The present and future of the Cancer Dependency Map. Nat Rev Cancer. Nature Research; 2025;25:59–73. 10.1038/s41568-024-00763-x

20. Yang M, Chitambar CR. Role of oxidative stress in the induction of metallothionein-2A and heme oxygenase-1 gene expression by the antineoplastic agent gallium nitrate in human lymphoma cells. Free Radic Biol Med. 2008;45:763–72. 10.1016/j.freeradbiomed.2008.05.031

21. Chitambar CR, Antholine WE. Iron-targeting antitumor activity of gallium compounds and novel insights into triapine®-metal complexes. Antioxid. Redox Signal. 2013. p. 956–72. 10.1089/ars.2012.4880

22. Meng EC, Goddard TD, Pettersen EF, Couch GS, Pearson ZJ, Morris JH, et al. UCSF ChimeraX: Tools for structure building and analysis. Protein Science. John Wiley and Sons Inc; 2023;32. 10.1002/pro.4792

23. Li F, Liu F, Huang K, Yang S. Advancement of Gallium and Gallium-Based Compounds as Antimicrobial Agents. Front Bioeng Biotechnol. 2022;10:1–15. 10.3389/fbioe.2022.827960

24. Bernstein LR, Park M. Gallium, Therapeutic Effects. Encyclopedia of Metalloproteins. 2013; 10.1007/978-1-4614-1533-6

25. Sun W, Qi M, Cheng S, Li C, Dong B, Wang L. Gallium and gallium compounds: New insights into the “Trojan horse” strategy in medical applications. Mater Des [Internet]. The Authors; 2023;227:111704. 10.1016/j.matdes.2023.111704

26. Chitambar CR. Gallium and its competing roles with iron in biological systems. Biochim Biophys Acta Mol Cell Res [Internet]. Elsevier B.V.; 2016;1863:2044–53. 10.1016/j.bbamcr.2016.04.027

27. Lane AN, Fan TWM. Regulation of mammalian nucleotide metabolism and biosynthesis. Nucleic Acids Res. Oxford University Press; 2015. p. 2466–85. 10.1093/nar/gkv047

28. Popović-Bijelić A, Kowol CR, Lind MES, Luo J, Himo F, Enyedy ÉA, et al. Ribonucleotide reductase inhibition by metal complexes of Triapine (3-aminopyridine-2-carboxaldehyde thiosemicarbazone): A combined experimental and theoretical study. J Inorg Biochem. 2011;105:1422–31. 10.1016/j.jinorgbio.2011.07.003

29. Casey Zoss RT, Nd, YS, Ks, CC. Ferroportin-dependent iron efflux as a novel mechanism of gallium maltolate in glioblastoma. 10.1093/neuonc/noaf201.0698

30. Pang Z, Zhou G, Ewald J, Chang L, Hacariz O, Basu N, et al. Using MetaboAnalyst 5.0 for LC– HRMS spectra processing, multi-omics integration and covariate adjustment of global metabolomics data. Nat Protoc. Springer US; 2022;17:1735–61. 10.1038/s41596-022-00710-w

