## Supplemental information for "Gallium induces cytotoxicity through disruption of DNA synthesis rather than ferroptosis"

### Supplementary Figures 1-5

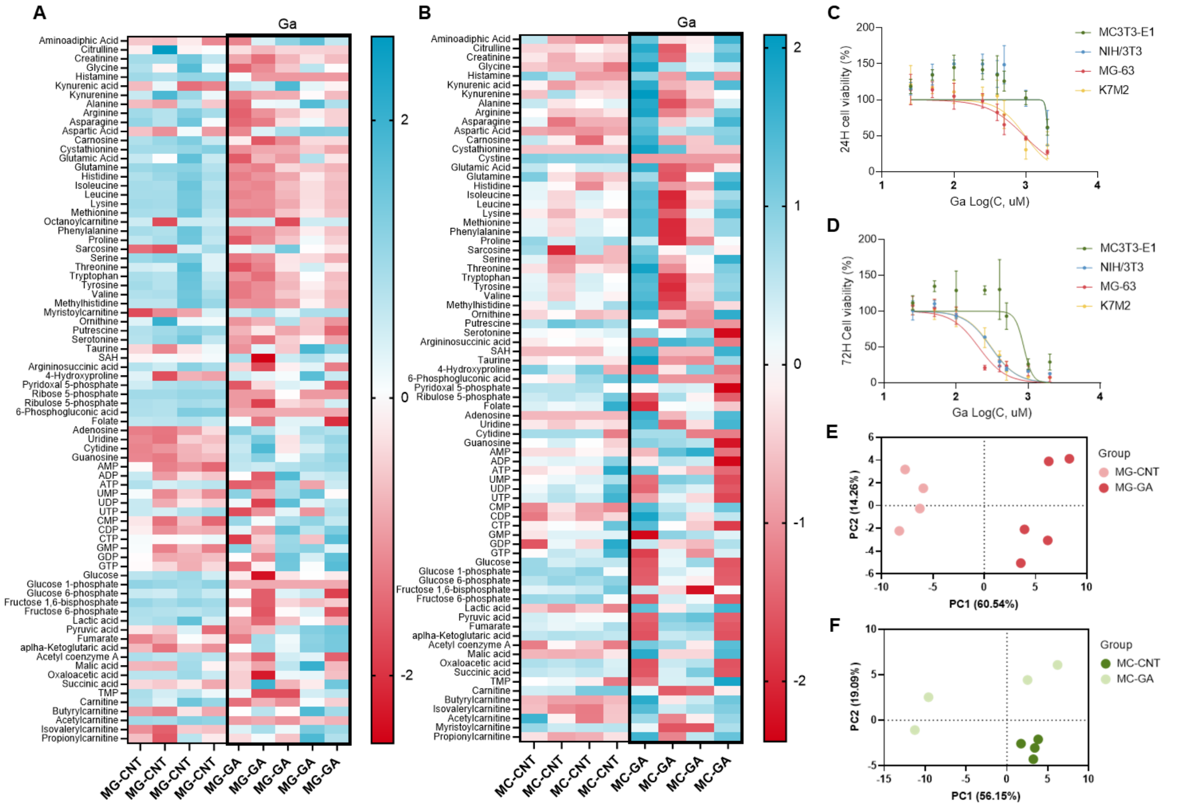


**Figure S1. Overall view of metabolomics results and IC50 to both bone cell lines. Related to Figure 1. (A-B) Heatmap of metabolomic results**, A for MG-63 and B for MC3T3-E1. Metabolites are grouped as nucleosides, sugar metabolites, carnitine family, and amino acids. **(C-D)** The IC50 of gallium to MC3T3-E1 was 2039 μM (24 hours) and 869.4 μM (72 hours). The IC50 of gallium to MG-63 were 948.4 μM (24 hours) and 205.2 μM (72 hours). The IC50 of gallium to NIH/3T3 was 2036 μM (24 hours) and 302.3 μM (72 hours). The IC50 of gallium to K7M2 was 975.4 μM (24 hours) and 308.7 μM (72 hours). **(E-F)** PCA plots of the metabolomic results, **E** for MG-63 and **F** for MC-3T3.

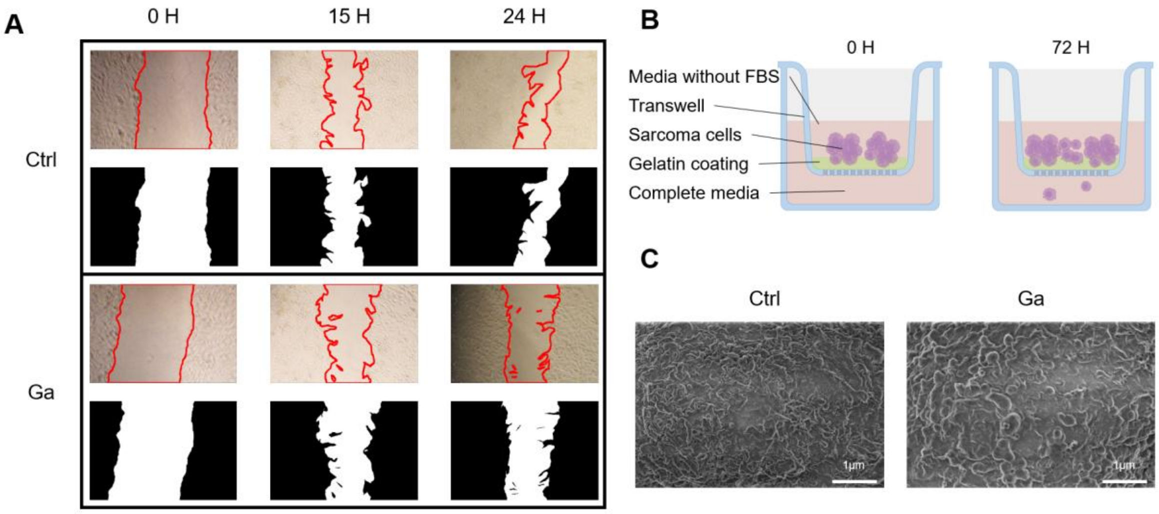

**Figure S2. Gallium disrupts the cytoskeletal structure and blocks sarcoma migration. Related to Figure 2. (A) Cell migration assay images: original (top) and contrast-enhanced (bottom).** Numerous rounded cells are evident in the gallium-treated groups at 15 h and 24 h, indicating gallium-induced changes in cell morphology. **(B) Schematic of the Transwell invasion assay (created with Figdraw).** The upper chamber contained serum-free medium; the lower chamber contained 10% FBS as a chemoattractant. **(C) The SEM images of cell membrane.** No rupture or holes in the cell membrane were found.

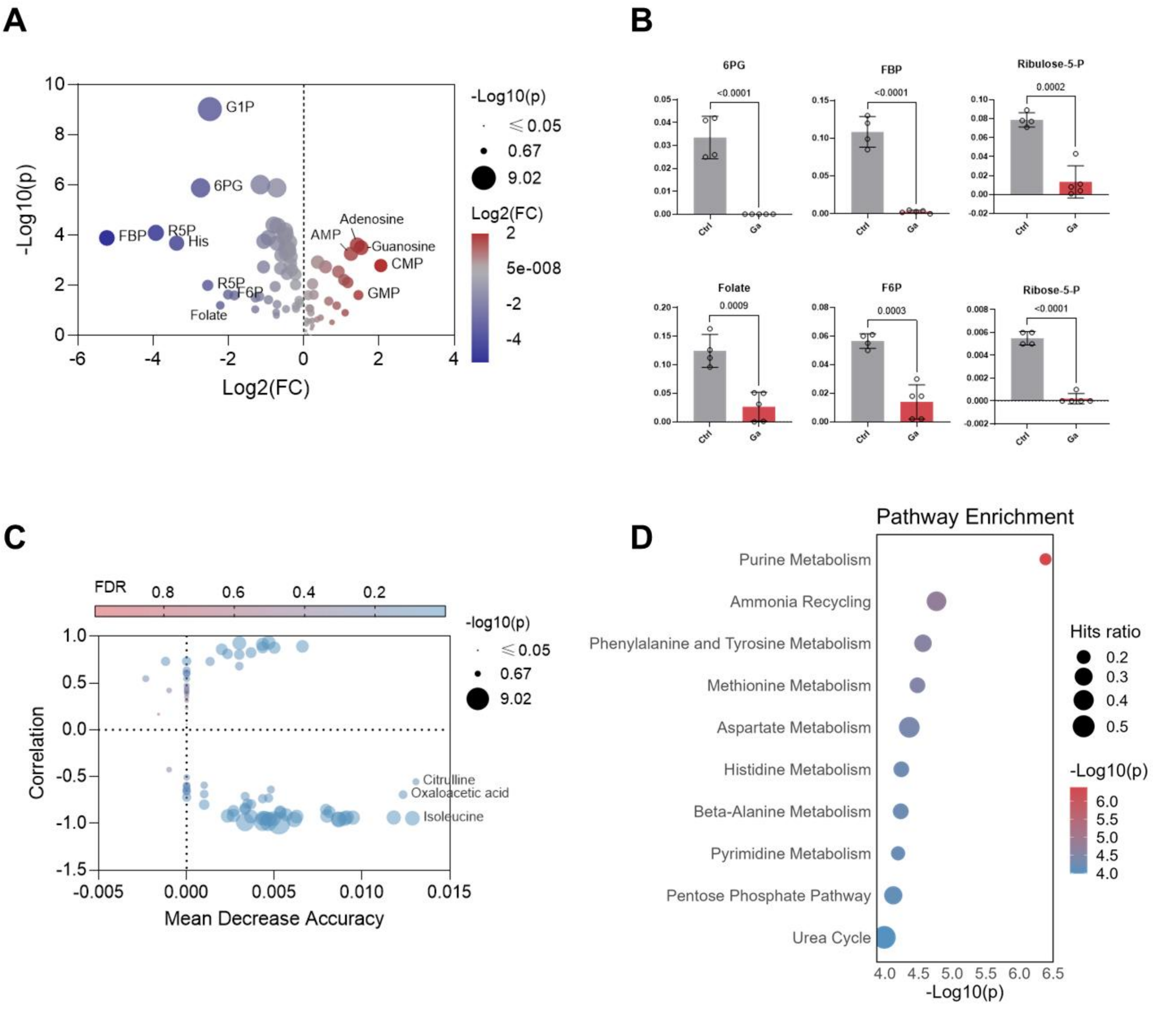


**Figure S3 Supplementary figure of metabolomics of gallium-treated MG-63 cells. Related to Figure 3. (A) Volcano plot of differential metabolites. (B) Concentrations of selected cellular metabolites in MG-63 cells referenced in the statistical analysis (y-axis: µM). (C) Integrated feature ranking combining correlation pattern search (correlation score) and Random Forest (MDA).** Both Correlation pattern search Random Forest were conducted based on treated/not-treated with gallium. **(D) Pathway enrichment analysis based on Human Metabolome Database (HMDB).**

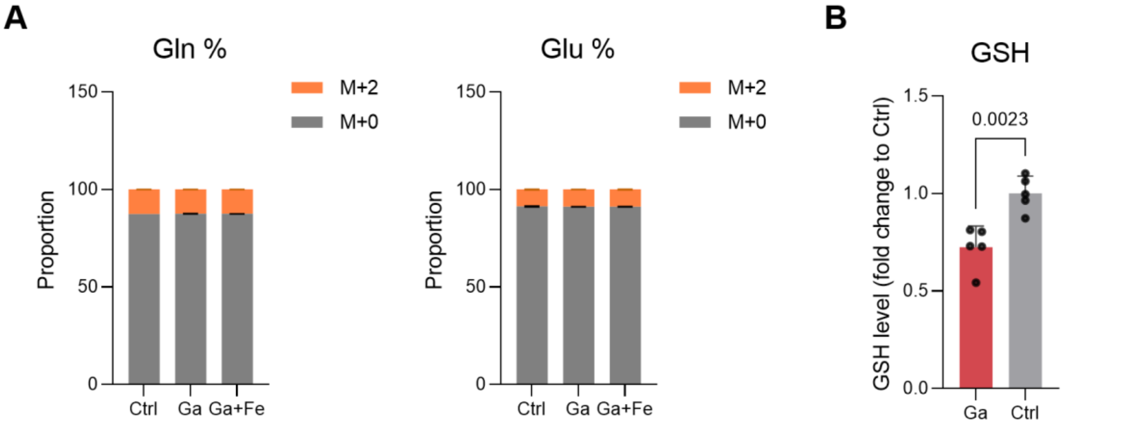


**Figure S4 Supplementary figure of isotope tracing. Related to Figure 4. (A)** The isotopic ratios of glutamine and glutamate are similar in each group. **(B)** The GSH level of MG-63 cells treated by gallium compared with the control, conducted by GSH assay kits, unpaired t-test.

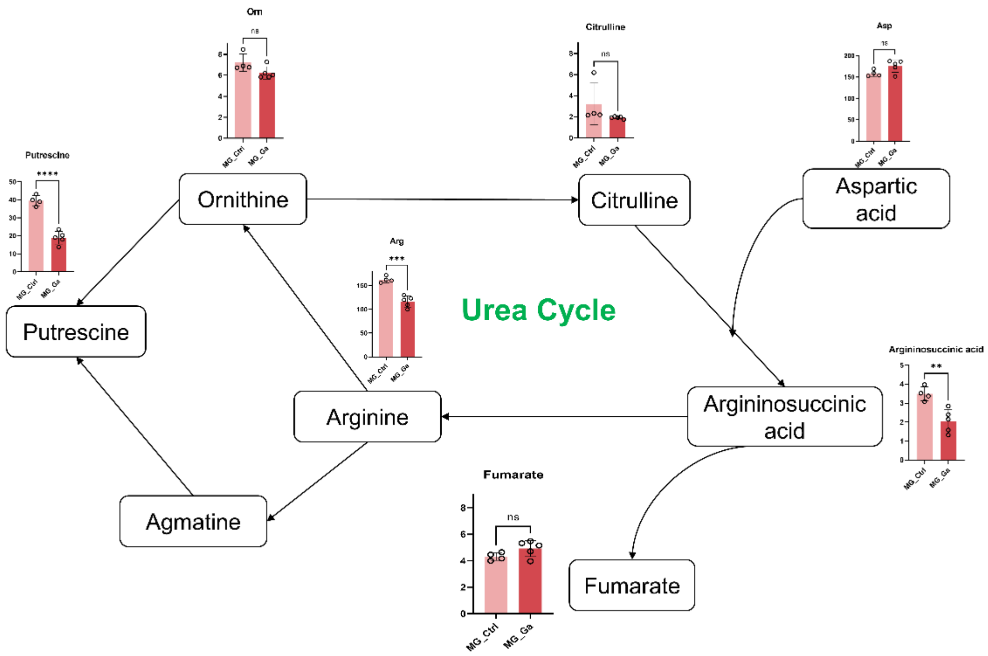


**Figure S5 Urea cycle metabolite changes of MG-63 under gallium treatment.** Provided as supplementary material for readers' reference.

### Supplementary Tables 1-3

| Ion | Charge | Coordination | Spin State | Crystal Radius | Ionic Radius |
|---|---|---|---|---|---|
| Fe | 2 | IV | High Spin | 0.77 | 0.63 |
| | | IVSQ | High Spin | 0.78 | 0.64 |
| | | VI | High Spin | 0.92 | 0.78 |
| | | | Low Spin | 0.75 | 0.61 |
| | | VIII | High Spin | 1.06 | 0.92 |
| | 3 | IV | High Spin | 0.63 | 0.49 |
| | | V | | 0.72 | 0.58 |
| | | VI | Low Spin | 0.69 | 0.55 |
| | | | High Spin | 0.785 | 0.645 |
| | | VIII | High Spin | 0.92 | 0.78 |
| | 4 | VI | | 0.725 | 0.585 |
| | 6 | IV | | 0.39 | 0.25 |
| Ga | 3 | IV | | 0.61 | 0.47 |
| | | V | | 0.69 | 0.55 |
| | | VI | | 0.76 | 0.62 |
| Cu | 1 | II | | 0.6 | 0.46 |
| | | IV | | 0.74 | 0.6 |
| | | VI | | 0.91 | 0.77 |
| | 2 | IV | | 0.71 | 0.57 |
| | | IVSQ | | 0.71 | 0.57 |
| | | V | | 0.79 | 0.65 |
| | | VI | | 0.87 | 0.73 |
| | 3 | VI | Low Spin | 0.68 | 0.54 |

**Table S1. A Comparison of Radii for Fe, Ga, and Cu**(1)

| **Protein** | **Function** | **Subcellular location** |
|---|---|---|
| Respiratory complexes I–III | Electron transport chain | Mitochondrial inner membrane |
| Mitochondrial aconitase | TCA cycle | Mitochondrial matrix |
| Ferrochelatase | Haem biosynthesis | Mitochondrial matrix |
| NTH1/MYH | DNA repair | Mitochondrial matrix, nucleus |
| IRP1/cytosolic aconitase | Iron metabolism | Cytosol |
| Xanthine oxidase | Purine metabolism | Cytosol |
| GPAT (ATase) | Purine biosynthesis | Cytosol |
| Methionine synthase | Methionine biosynthesis | Mitochondrial, cytosol |
| Ribonucleotide reductase | DNA synthesis and repair | Cytosol, nucleus |
| Catalase | breakdown of H2O2 | Peroxisomes |

**Table S2. Proteins that functionalized with iron**

| Pathway | Direction | Key Metabolites Affected |
|---|---|---|
| Glycolysis(2) | Down (↓) | Lactate, Glucose-6-Phosphate, Pyruvate |
| TCA Cycle(2) | Down (↓) | Citrate, Malate, Succinate, ATP |
| Redox/Detox(3) | Mixed | ↓ GSH (depleted), ↑ GSSG (oxidized form) |
| Amino Acids(4) | Up (↑) | Proline, Methionine metabolites, Phenylalanine |
| Nucleotides(4,5) | Down (↓) | PRPP, Purine/Pyrimidine nucleotides |
| Lipids(6) | Up (↑) | Ceramides (death signal), Lipid Droplets |

**Table S3. Metabolites change of cells under Cis-platin treatment**

### References


1. Shannon RD. Revised effective ionic radii and systematic studies of interatomic distances in halides and chalcogenides. Acta Crystallographica. 1976;32(5):751–67.

2. Guo J, Yu J, Peng F, Li J, Tan Z, Chen Y, et al. In vitro and in vivo analysis of metabolites involved in the TCA cycle and glutamine metabolism associated with cisplatin resistance in human lung cancer. Expert Rev Proteomics. 2021;18(3):233–40.

3. Dong XQ, Chu LK, Cao X, Xiong QW, Mao YM, Chen CH, et al. Glutathione metabolism rewiring protects renal tubule cells against cisplatin-induced apoptosis and ferroptosis. Redox Report. 2023;28(1).

4. Wahwah N, Dhar D, Chen H, Zhuang S, Chan A, Casteel DE, et al. Metabolic interaction between amino acid deprivation and cisplatin synergistically reduces phosphoribosyl-pyrophosphate and augments cisplatin cytotoxicity. Sci Rep. 2020 Dec 1;10(1).

5. Hu J, Lieb JD, Sancar A, Adar S. Cisplatin DNA damage and repair maps of the human genome at single-nucleotide resolution. Proc Natl Acad Sci U S A. 2016 Oct 11;113(41):11507–12.

6. Dupre T V., Doll MA, Shah PP, Sharp CN, Siow D, Megyesi J, et al. Inhibiting glucosylceramide synthase exacerbates cisplatin-induced acute kidney injury. J Lipid Res. 2017;58(7):1439–52.
